# Evolution of virulence in emerging epidemics: inference from an evolution experiment

**DOI:** 10.1101/2020.08.19.256917

**Authors:** François Blanquart, Thomas Berngruber, Marc Choisy, Sylvain Gandon

## Abstract

Inference using mathematical models of infectious disease dynamics is a powerful tool to analyse epidemiological data and elucidate pathogen life cycles. Key epidemiological parameters can be estimated from demographic time series by computing the likelihood of alternative models of pathogen transmission. Here we use this inference approach to analyze data from an evolution experiment in which we monitored both the epidemiology and the evolution of the temperate bacteriophage *λ* during an epidemic. We estimate parameter values for all the life-history traits of two distinct strains of the virus. In particular, we estimate the ability of the two virus strains to modulate plastically the rate of lysogenization with the multiplicity of infection. Our work illustrates how inference from experimental evolution data can feedback on the development of models aiming to predict the epidemiology and evolution of infectious diseases.

## Introduction

Evolutionary epidemiology theory is a framework that aims to capture both the short-term and long-term evolutionary dynamics of pathogens (1–3). It provides a way to bridge the gap between mathematical epidemiology and population genetics and shows how the evolution of pathogen virulence is driven by the change in the density of susceptible hosts. At the onset of an epidemic, when the incidence of the disease is small, evolutionary theory predicts that selection for pathogen transmission is strongest because there are many opportunities to infect new hosts. If there is a positive genetic covariance between the rate of transmission and the virulence of the pathogen, one may expect to see selection for pathogen virulence at the beginning of an epidemic. Later on, the spread of the infection reduces the pool of susceptible hosts and may reverse the selection on virulence (1–5).

We explored the validity of this prediction with an experimental approach where we studied the evolution of bacteriophage *λ* throughout the development of an epidemic in continuous cultures of *Escherichia coli* (6). Bacteriophage *λ* is a typical temperate virus which integrates into the host genome and transmits vertically to daughter cells at cell division. Integration of phage *λ* into the genome protects the host cell against superinfection of other *λ* phage particles and this way provides immunity to superinfection by other *λ* phage particles (7). The reactivation of the integrated phage (the prophage) results in lysis and destruction of the host cell. Lysis of its host prevents vertical transmission but allows the phage to be transmitted horizontally to uninfected susceptible cells. Whereas the non-virulent *λ* wildtype transmits mostly vertically by dormant integration into the host genome, the virulent mutant *λ*cI857 transmits mostly horizontally by host lysis. This difference in virulence and transmission mode is the result of a point mutation in the *λ* virulence repressor protein cI which actively controls the decision to lyse the host cell. We monitored the competition of the bacteriophage *λ* and its virulent mutant *λ*cI857 and compared these dynamics with the predictions from a mathematical model that tracks both the epidemiology and the evolution of the virus (6). The parameter values used for the model where obtained from independent estimates in previous studies and were used for an experimental validation of the theoretical predictions from our model.

In the present work, we go beyond the qualitative comparison between the theory and the experiments. We use the experimental data on the prevalence of infection and on the frequency of the virulent mutant to fit a dynamical model describing the epidemiology and the evolution of the phage. This approach provides a rigorous way to quantify the phenotypic differences between the two virus strains used in our experiments and in particular quantify key life-history traits like the probabilities of integration and the rates of reactivation. This approach also provides a way to compare the likelihoods of alternative models of virus transmission (with or without phenotypic plasticity).

## Methods

### An evolutionary epidemiology model

Following Berngruber *et al.* (6) we model the dynamics of the bacterial population infected by a polymorphic population of phage *λ* in a chemostat. The density of susceptible (uninfected) bacteria is called *S*. The two strains of phage, called the “wild type” (WT) and “mutant” (M), have a low and a high probability to cause a lytic (rather than lysogenic infection), respectively. The density of bacteria infected by the strain *i* (*i* ∈ {*WT*, *M*}) in a lysogenic state is *L*_*i*_, and *L* = ∑_*i*_ *L*_*i*_ is the total density of lysogens. The density of free viral particles of type *i* is *V*_*i*_, and *V* = ∑_*i*_ *V*_*i*_ is the total density of free virus. In the absence of infection, bacteria reproduce at a density-dependent rate 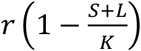, where *K* is the carrying capacity of the bacteria. Infected (lysogenized) bacteria are assumed to reproduce at the same rate as susceptible bacteria. The outflow of the chemostat removes bacteria and free phages at a rate *m*. Infection occurs when free viruses adsorb to a susceptible cell (at a rate *a*) and enters the cell (with a probability *b*). The rate of infections is thus *abV*. Infection by phage strain *i* gives rise either to a lysogenic bacterium, with probability *ϕ*_*i*_, or may immediately lyse the bacterium and release a number *B* of free phages, with probability 1 – *ϕ*_*i*_. The parameter *ϕ*_*i*_ determines the probability of genome integration of the phage. The variable *ϕ*_•_ is the mean probability of integration of free phages defined by:

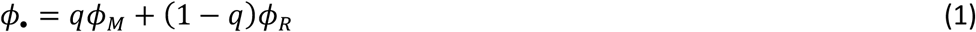

where *q* is the frequency of mutant phage among free viruses. Consequently, new lysogenic bacteria are created at a rate *abϕ*_•_*VS*, while new free phages are created at a rate *ab*(1 − *ϕ*_•_)*VSB*.

The prophage in lysogenic bacteria may switch to a lytic cycle, with rate *α*_*i*_. The induction of the lytic cycle kills the lysogenic bacteria and yields *B* free phages of type *i*. The variable *α*_∘_ is the mean rate of reactivation of viruses in latently infected bacteria, defined as:

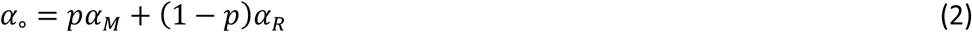

where *p* is the frequency of mutant phage among prophages.

The above life cycle yields the following system of ordinary differential equations describing the epidemiological dynamics (the change in the total densities of different bacteria and the total density of free phages), and the evolutionary dynamics (the change in the frequencies of the mutant phage in the prophage and the free virus compartment):

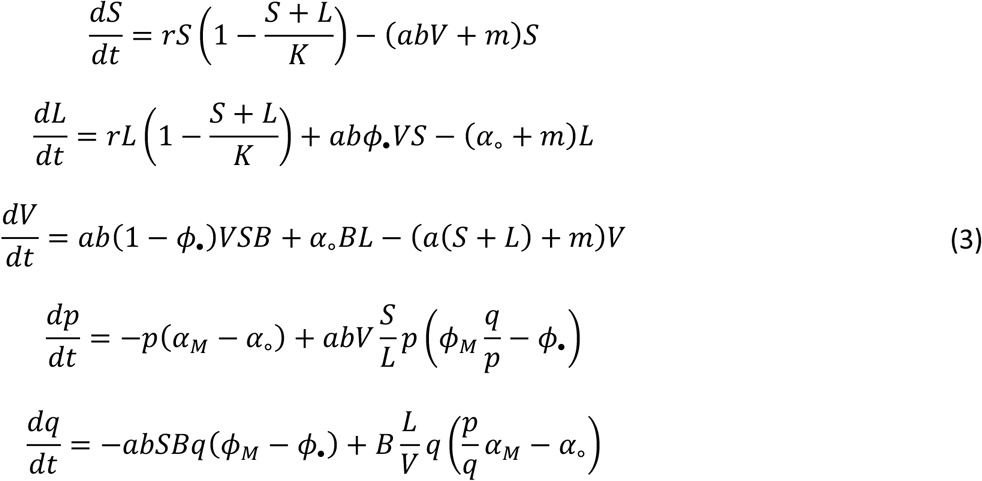

### Fitting the model to experimental data from a chemostat experiment

We take a maximum likelihood approach to fit the above model to different types of data obtained from the experiment. First, we use the prevalence of the phage infection which refers to the proportion of lysogenic cells:

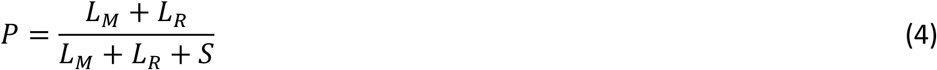

where the relative densities of the different types of cells where obtained by flow cytometry. Second, we used the flow cytometry data to obtain the frequency of mutant phage among lysogenic bacteria:

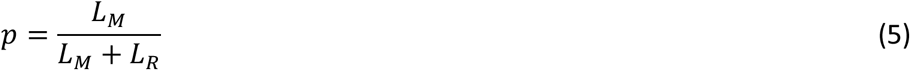

Third, we used quantitative PCR to measure the relative densities of wild type and mutant free viruses and to obtain the frequency of free mutant phages *q*:

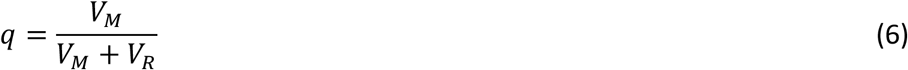

These three variables (prevalence, frequency of mutant phage in infected cells, frequency of free mutant phages) were measured in four replicate chemostats for a treatment with high initial phage prevalence—the *endemic* treatment—and four replicate chemostats for a treatment with low initial phage prevalence—the *epidemic* treatment. The chemostat dynamics were followed approximately every hour for a maximum of 60 hours.

The likelihood of this data is the probability of the data given the dynamical model. As there is no closed-form analytical solution for the model, we integrated it numerically. We assumed that the logit-transformed prevalence and frequencies are measured with a normally distributed error. The likelihood corresponding to the observed prevalence in all treatments, replicates and time is defined as:

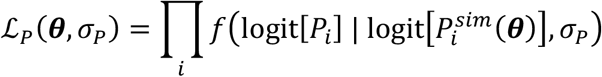

where *f*(*x* | *μ*, *σ*) is the density of a normal distribution with mean *μ* and standard deviation *σ*, evaluated at *x*. The product is over all observations, where the index *i* scans the two treatments, four replicates for each treatment and up to 60 time points for each replicate. *P*_*i*_ denotes the *i*th observation of prevalence. 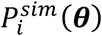 is the prevalence corresponding to observation *i* when integrating the ODE model with parameters *θ*. We use the logit-transformed prevalence: logit(*P*_*i*_)=log(*P*_*i*_/(1−*P*_*i*_)). The parameter *σ*_*P*_ is the standard deviation of the measurement error corresponding to the logit-transformed prevalence. It is assumed to be the same for all treatments, replicates and time points.

Similarly, the likelihoods corresponding to the mutant frequencies among infected bacteria and among free phages are:

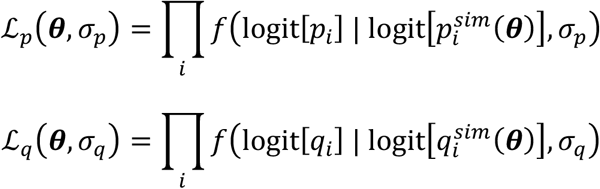

These likelihoods also depend on the set of parameters of the ODE model with parameters ***θ***. The parameters *σ*_*p*_ and *σ*_*q*_ refer to the standard deviation of the measurement error associated with *p* and *q*, respectively. Note that for the sake of simplicity, such noise is assumed to be uncorrelated across replicates, and across time points of the same replicate. We thus neglect the demographic stochasticity that may generate additional variation around the average dynamical process. This is justified by the large population sizes of both the bacteria and the phage in the chemostat. Finally, we defined an overall composite likelihood where the errors on prevalence and frequencies data are assumed to be independent between them, as:

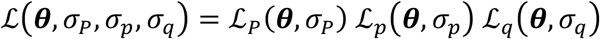

We fitted the model by maximum likelihood methods, maximizing the function 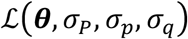 with respect to the parameters ***θ***, *σ*_*P*_, *σ*_*p*_, *σ*_*q*_, using the Nelder-Mead algorithm. To ensure convergence to a global optimum, we repeated each optimisation multiple times from random starting points and selected the overall maximum likelihood. We computed 95% confidence intervals for the parameters from MCMC sampling of the likelihood function.

The simpler model without plasticity being nested in a more complex model where the phage additionally exhibits plastic lysogeny, we compare the fit of the two mechanistic models using a likelihood ratio test.

Computations were done with R, using the deSolve package for numerical integration of differential equations (8,9).

## Results

### Maximum likelihood estimation

Of the 13 parameters of the model, 2 were fixed to the values chosen in the experimental design (6). Eleven parameters were estimated by maximum likelihood (**Table 1**), including eight parameters from the model and three error standard deviations. The maximum log-likelihood of this model was *L*_1_ = −1911.96. Many parameters could be estimated with narrow confidence intervals, an exception being the burst size which could be between 44 and 488 phages per cell. In other words, the dynamics of the three variables *P*, *p* and *q* contain information on most model parameters. When computing the dynamics with the maximum likelihood parameters, the dynamics of prevalence and frequency of resistance generally closely fitted the data (**Figure 1**). However, the frequency of bacteria infected by the mutant phage peaked sharply in the epidemic phase in the treatment with low initial prevalence, an observation that the model failed to reproduce.

**Table 1:**
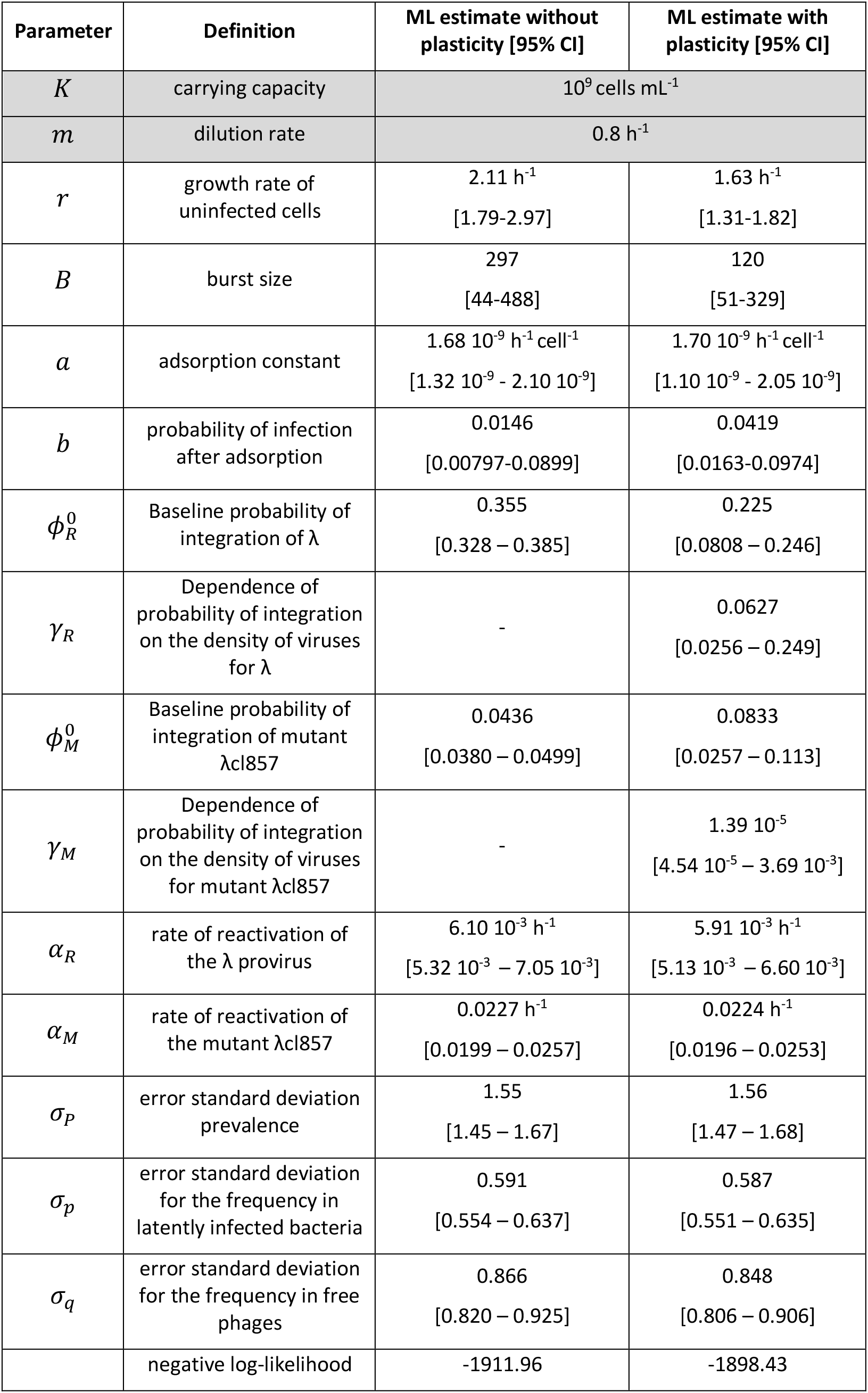
Model parameters. Two parameters (K and m) were fixed in the experiments (6). The 13 other parameters were estimated using the model without or with plasticity for the probability of lysogenisation. The table indicates the ML estimation and the 95% confidence interval. The bottom line of the table gives the maximum log-likelihood of the models without or with plasticity.

**Figure 1:**
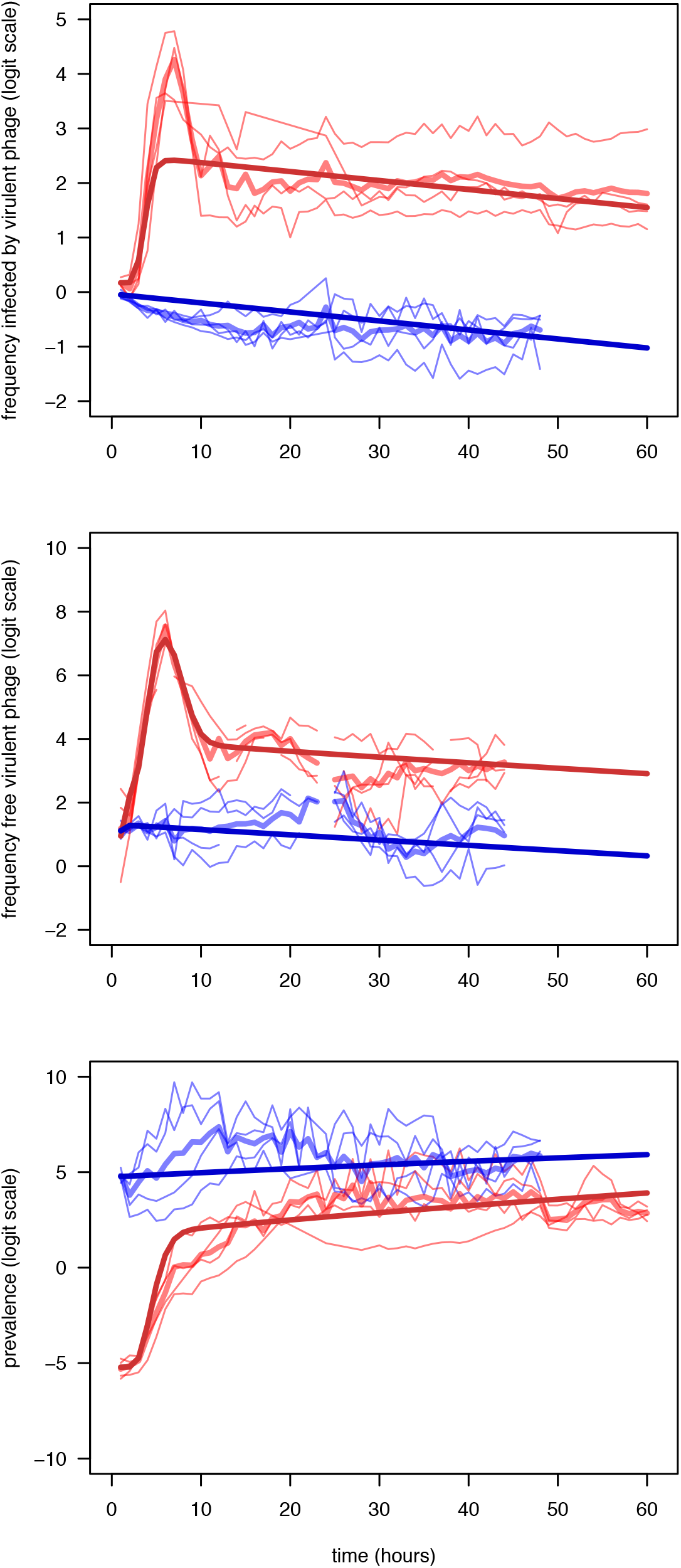
viral evolution in the chemostat, data (light blue and red lines) and model (dark blue and red thick lines). The blue curves represent the endemic conditions (high initial prevalence), and the red curves the epidemic conditions (low initial prevalence). We show the data for each replicate (light thin lines) as well as the mean across replicates (thick line). Only the predictions of the extended model with plasticity are shown. Top panel: the logit frequency of mutant (virulent) phage in lysogenic state in infected cells. Middle panel: the logit frequency of free mutant phage. Bottom panel: the logit prevalence of infection.

### Extended model including the plasticity of phage *λ*

The phage *λ* is able to modulate the probability of lysogenisation with the multiplicity of infection of the bacteria (10,11). We examined whether the experimental data was better explained by accounting for plasticity in lysogenisation. We extended the first model to additionally allow the probability of integration to respond plastically to the multiplicity of infection, which depends on the density of free viruses *V* (Supplementary Information)

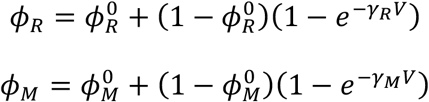

The parameter 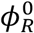 is the probability of integration of the wild type strain when a single phage infects a bacteria. The probability of lysogenisation increases with the density of viruses and the parameters *γ*_*R*_ and *γ*_*M*_ govern the magnitude of this effect for both the wild type and the mutant, respectively.

In the simulations of the best fit model (without plasticity), the density of viruses *V* varies from 0 to 1.4· 10^9^ viruses in the endemic conditions, and from 0 to 7· 10^10^ viruses in the epidemic conditions. We found that the model with plasticity had a statistically significant better support than the simpler model (*L*_2_ = −1898.43, *χ*^2^ = 27.06, *p* = 1. 10^−16^}). This model finds a strong plastic relationship between the density of virus and the probability of integration in the wild type strain of the virus, and no plasticity in the mutant strain (**Figure 2**). The model with plasticity yields a better fit to the dynamics of the frequency of the mutant phage during the epidemic in the free virus compartment (**Figure S1**), which explains its better likelihood.

**Figure 2:**
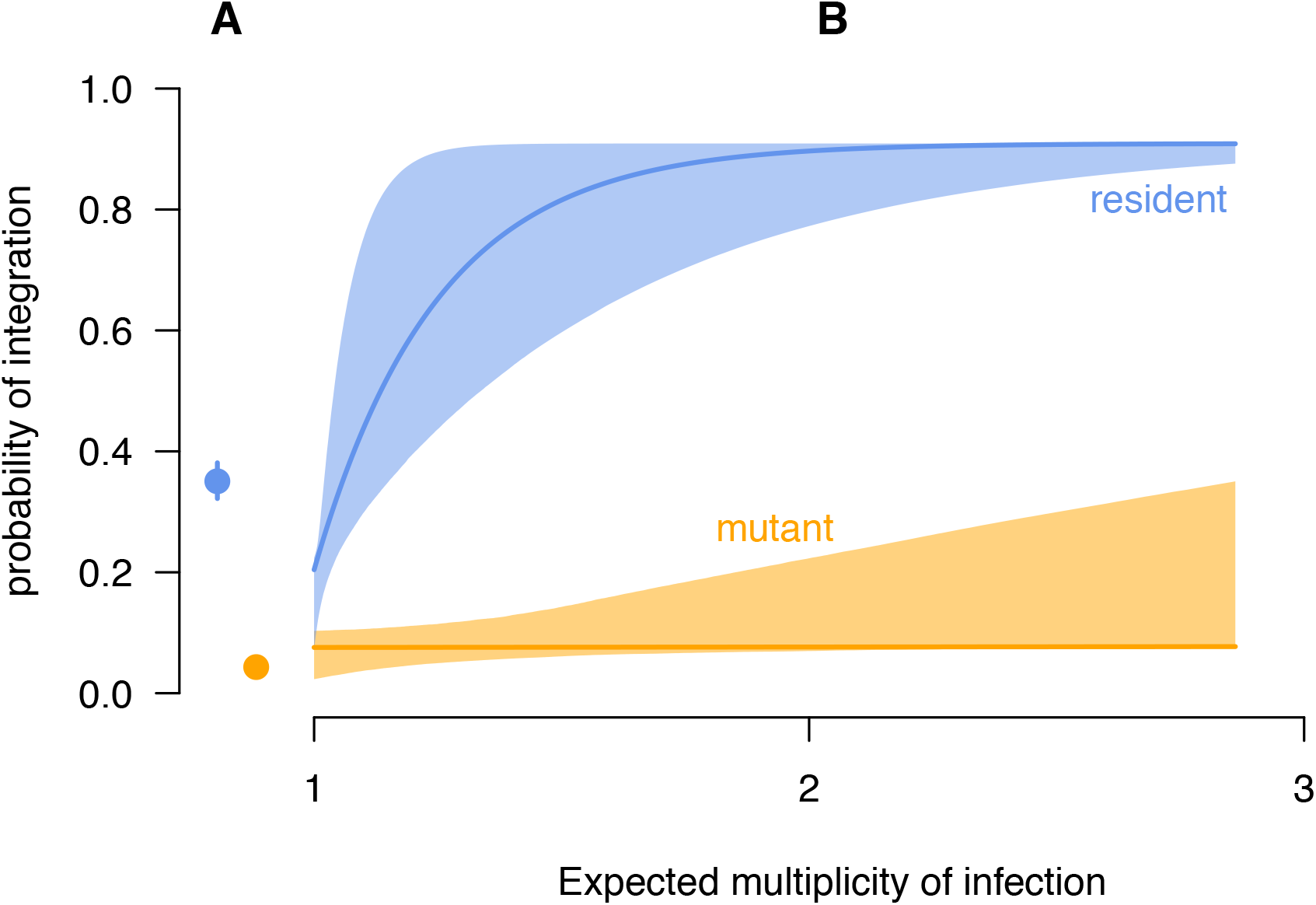
inferred relationship between the probability of integration and the expected multiplicity of infection (reflecting the density of free virus,), for the wild type phage λ (blue) and the mutant λcl857 (orange). Left part of the graph (A), the first model without plasticity; Right part of the graph (B), the second model with plasticity, where the probability of integration depends on the density of free viruses. Vertical segments (A) and shaded areas (B) represent 95% confidence intervals.

## Discussion

In a previous study we developed a theoretical model to study the evolution of a temperate virus during an epidemic (6). This model led to three main qualitative predictions: (i) The virulent mutant initially wins the competition with the wildtype when susceptible hosts are abundant, but the competitive outcome is reversed as soon as the epidemic reaches high prevalence; (ii) The virulent mutant is, at all times, more frequent among free viruses than among proviruses; and (iii) Lower initial prevalence results in a higher increase in frequency of the virulent mutant during the epidemic. In the same study we carried out an experiment with the bacteriophage *λ* which successfully tested the validity of these different predictions (6). The present work is an attempt to move in the opposite direction, from the experimental data back to the theoretical model.

First, under the assumption of the original model, we infer the parameter values that maximise the fit between the data and the simulated dynamics. This inference approach yields precise estimates for 8 different parameters (**Table 1**). All these estimations are consistent with previous estimates obtained from different studies (see Table S1 in Text S1 in (6)). This shows that combining data on the prevalence and on the frequency of different strains contains information on the specific values of the life-history traits (transmission, virulence) of different genotypes.

Second, we compared the original version of the model with a more general model that accounts for the observation that the phage can modulate its probability of lysogenisation with the multiplicity of infection determined by the density of free viruses. We explored this effect because we know that phage *λ* has the ability to adopt such plastic life-history strategies (10–12). The comparison of the two versions of the model reveals that the model with plasticity fits the data better than the original one. We also detect variations in plasticity between the wild type and the mutant phage (**Figure 2**). How does the estimated plastic response compare with the experimental measurement of the effect of multiple infections on the probability of lysogenisation by Zeng et al. (11)? Their measure of plasticity is based on the ability to track with time-lapse microscopy the fate (lysis versus lysogeny) of single bacteria that are exposed to 1, 2, 3, … phages. They found that the probability of lysogenisation increases from about 0.3 to 0.6 when the multiplicity of infection changes from 1 to 5. Our version of the model does not track the fate of individual cells and their multiplicity of infection. However, our model can be derived as an approximation of a more general model that explicitly describes the dynamics of cells exposed to a variable density of viruses (Supplementary Information). This more general model implies that the density of cells exposed to *i* viruses is geometrically distributed with parameter 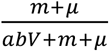, and the expected multiplicity of infection is 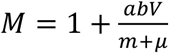. We could thus represent the probability of integration as a function of the expected multiplicity of infection (**Figure 2**). In our model, even in epidemic conditions (V = 6 × 10^10^), the expected multiplicity of infection is modest (around 2, Table S1) so most cells are theoretically simultaneously exposed to 1 or 2 phages. The inferred probability of integration is around 0.3 at low multiplicity of infection, which is consistent with the values measured by Zeng et al. However, our inferred relationship approaches 0.9 when the expected multiplicity of infection is 2, which is much higher than that measured experimentally, around 0.5. This difference between our population-based estimate and Zeng et al.’s result could be explained by differences in experimental conditions (e.g. temperature, liquid versus solid medium).

The plasticity of the wild type virus is a very important feature of the bacteriophage life cycle. In contrast, the mutant virus does not seem to exhibit plasticity. This is surprising because the original experiment by Kourilsky who monitored precisely the effect of multiplicity of infection on lysogenisation was performed with the *λCI*857 mutant. Note, however, that we carried out our experiments at 35°C while the experiments of Kourilsky were done at 32°C. The higher the temperature, the lower the rate of lysogenisation. The shape of the plasticity function could possibly be flattened by the increase in temperature (because of the destabilisation of the cI857 protein binding to the lambda operator (7). It would be interesting to explore this hypothesis using the experimental approach of Zeng et al. (11).

The present work allows to improve the theoretical model that we originally developed to describe the epidemiology and evolution of a temperate virus during an epidemic. This improved model can be used again to generate new predictions. For instance, we can use this epidemiological model of a temperate phage to explore the relationship between the density of free viruses and the density of cells in the environment. Several empirical studies have shown that the density of free viruses saturate when the density of bacteria increase in aquatic communities (13,14). What drives the relationship between host and free virus densities has led to some controversy (15–19). Our model can be used to explore the influence of virus plasticity on this relationship (using the two models defined in **Table 1**). **Figure 3** shows this relationship with or without plasticity when we assume a periodic fluctuation in the density of susceptible bacteria in the environment (see **Supplementary Information**). Phage plasticity increases the probability of integration when the multiplicity of infection increases, which reduces the release of free viruses in the environment. This mechanism, if it is widespread among aquatic viruses, could have a major influence on the functioning of marine ecosystems.

**Figure 3:**
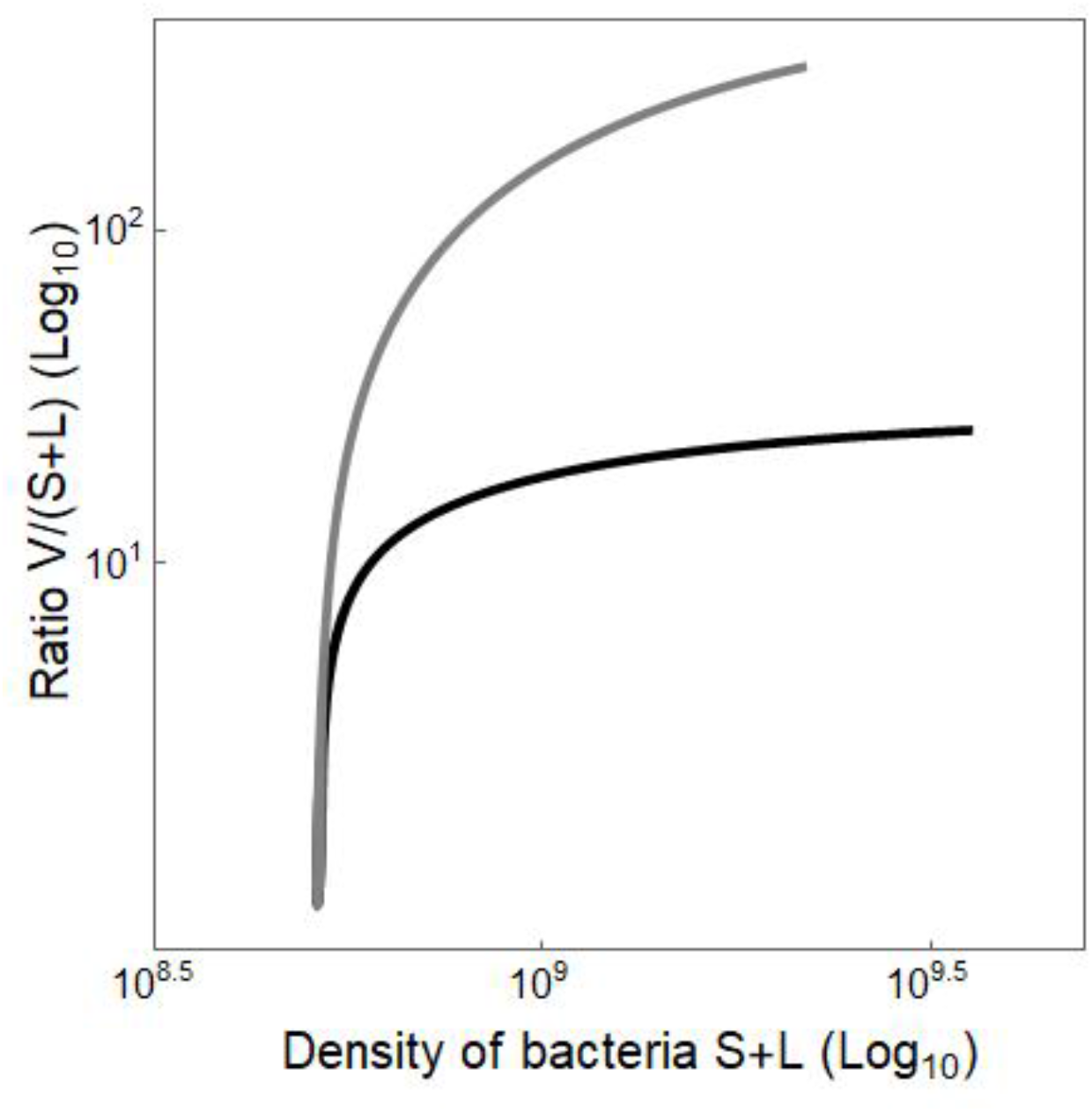
Relationship between the ratio of density of free virus and the density of bacterial cells in the environment for the two models described in Table 1: without plasticity (gray) and with plasticity (black).

In conclusion, statistical inference based on mechanistic models of infectious diseases can be a major tool to improve our understanding of the microbial dynamics monitored in evolution experiments. In fact, the back and forth between experiments and dynamical models may help design better experimental protocols were the aim will not only be to test a qualitative prediction but also to generate the data that will allow (1) the estimation of biologically relevant parameters and (2) the comparison of alternative dynamical models. This iterative process between model prediction and experimental validation is yielding a virtuous circle that improves the predictive power of evolutionary epidemiology theory.

## Supplementary information

### 1. Dynamics of multiple infections

We detail below a modified version of our model to account explicitly for the density *E*_*i*_ of bacterial cells exposed to *i* viruses. For the sake of simplicity, we only consider the simple case where the virus population is monomorphic (no mutant strain). This yields the following dynamical system:

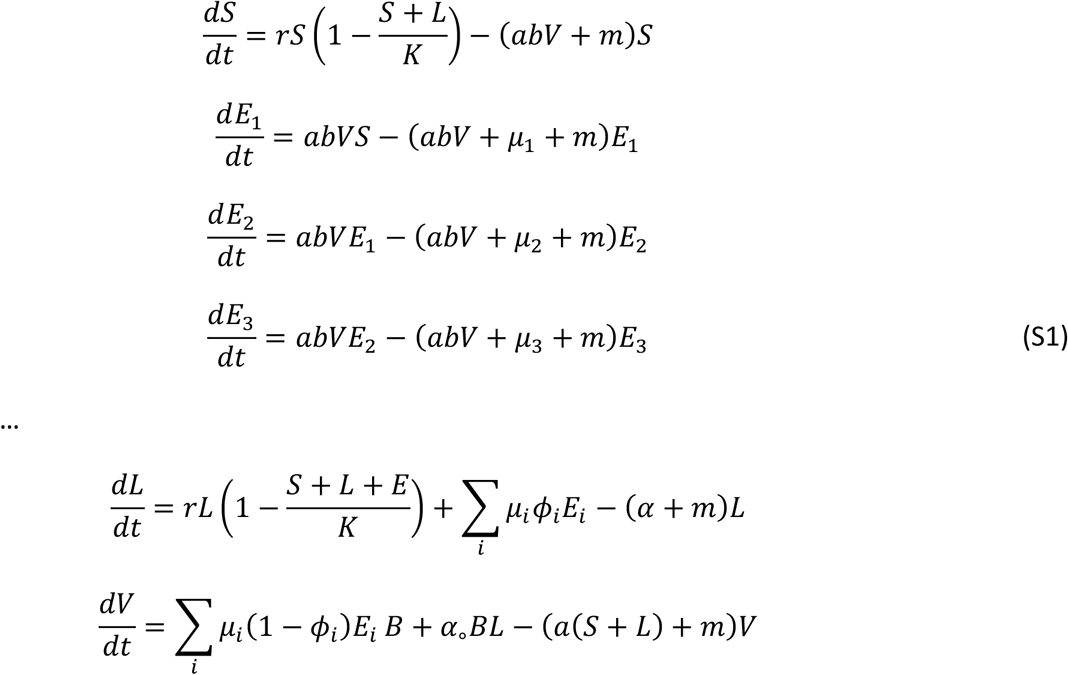

where *μ*_*i*_ is the rate at which a cell exposed to *i* viruses becomes infected (and thus turns into a lysogenic cell, or is lysed and liberates viruses). If we assume that the rates *μ*_*i*_ are relatively large we can use a separation of time scale argument to derive the quasi-equilibrium density of the different types of exposed cells:

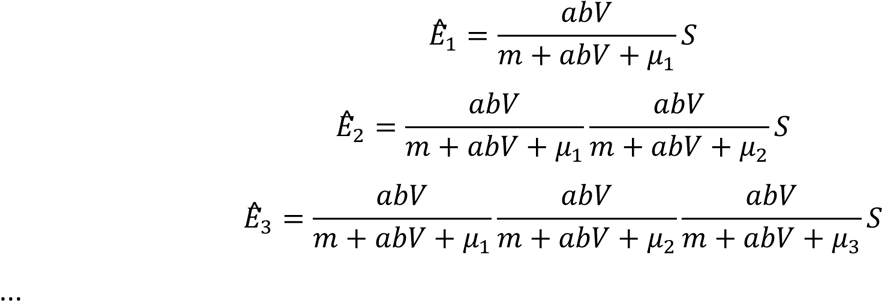

We can simplify this system using the additional approximation *μ*_*i*_ ≈ *μ*∀*i*, which yields:

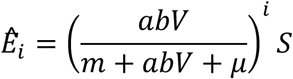

The frequency of cells exposed to *i* viruses is:

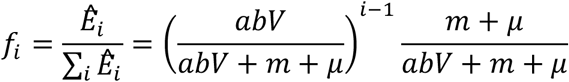

and the expected multiplicity of infection is:

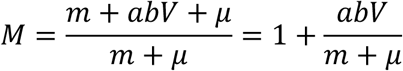

In other words, the frequency of cells exposed to *i* viruses follows a geometric distribution with parameter 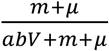.

This yields the following dynamical system:

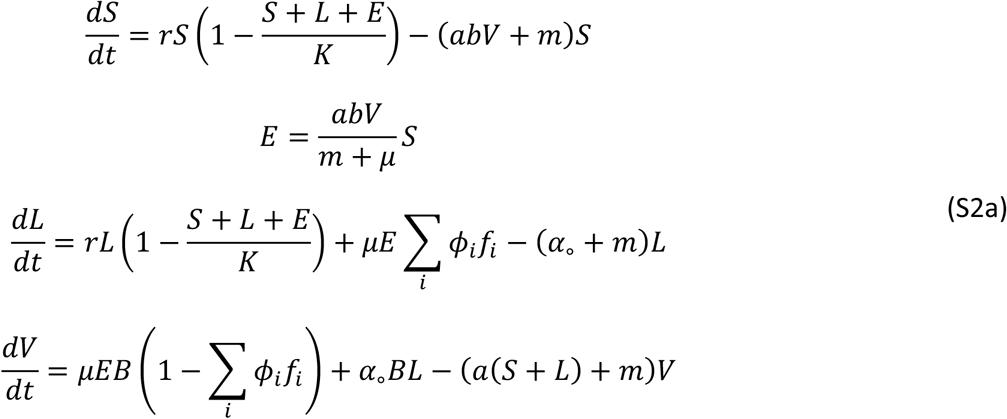

This system of equations is similar to the system defined in equation (3). The terms that differ are highlighted in blue. Note that when the rate at which an exposed cell becomes infected is large compared to the rate of exposition, and the rate of dilution (*μ* ≫ *abVS* and *μ* ≫ *m*), we recover the original model (equation (3) in the main text). Indeed, in that case, *E* is small and *μE* is approximately *abVS*, and the system becomes:

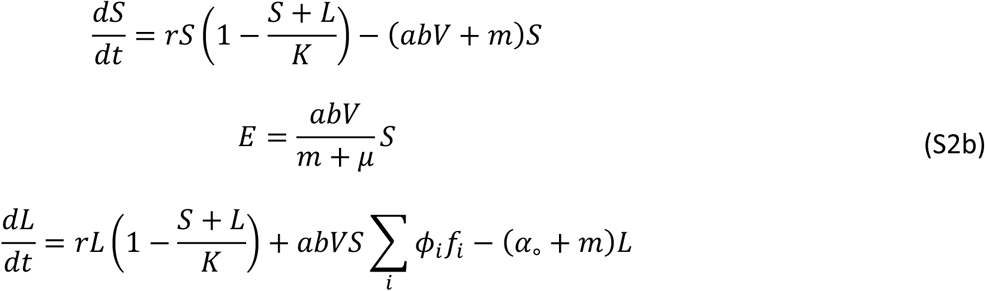

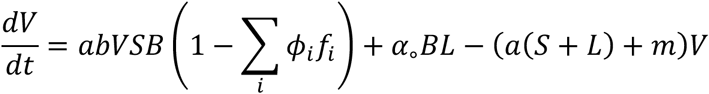

That model is analogous to the system in equation (3), except that here for simplicity the virus population is monomorphic. The mean probability of integration in that model is weighted by the frequencies of cells exposed to *i* virus, *f*_*i*_.

This model is particularly useful to estimate what is the expected multiplicity of infection *M* in our dynamical system. The multiplicity of infection depends on the density of free viruses *V*. We estimate here the multiplicity of infection at the endemic state and at the epidemic peak, where *V* is estimated to be 10^9^ and 6 × 10^10^ viruses per mL respectively. We estimate the rate at which an exposed cell becomes infected at *μ* = 1.5*h*^−1^. Indeed, in an experimental study (Zeng et al), single cells were observed for two hours to determine the outcome of infection (lysis or lysogeny). Assuming that the fate of 95% of exposed cells is decided in two hours gives a mean time to infection of 40 minutes, i.e. *μ* = 1.5*h*^−1^.

**Table S1:**
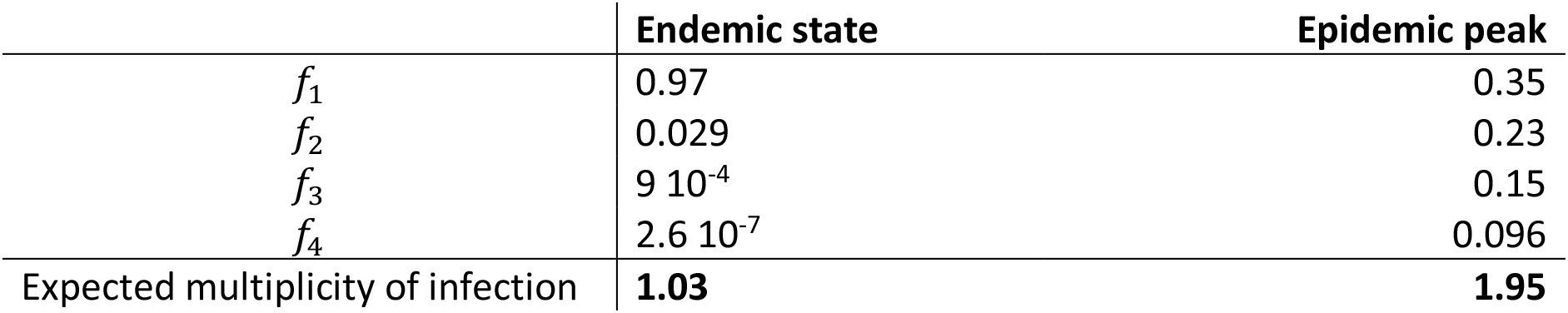
the estimated frequency of cells exposed to 1, 2, 3, 4 viruses, and the estimated expected multiplicity of infection when the system is at endemic state (V = 10^9^) and the system is at the epidemic peak (V = 6 10^10^).

|  | Endemic state | Epidemic peak |
| --- | --- | --- |
| $f_1$ | 0.97 | 0.35 |
| $f_2$ | 0.029 | 0.23 |
| $f_3$ | $9 \cdot 10^{-4}$ | 0.15 |
| $f_4$ | $2.6 \cdot 10^{-7}$ | 0.096 |
| Expected multiplicity of infection | **1.03** | **1.95** |

### 2. Dynamics of the ratio *V*/(*S* + *L*) in temporally variable environments

To explore the influence of phage plasticity in a variable environment we use the following model:

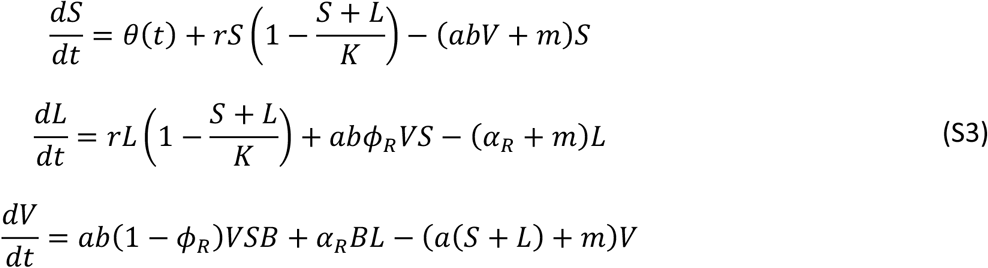

where 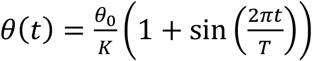 is a function which is meant to induce a periodic fluctuation (seasonal) of the abundance of susceptible cells. These fluctuations result in periodic variations of the densities of free virus and bacterial cells. **Figure 3** in the main text plots the relationship between 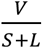 and *S* + *L* when the system has reached a periodic equilibrium. Parameter values: *θ*_0_ = 10, *T* = 10000*h*. Other parameter values as in Table 1.

**Figure S1:**
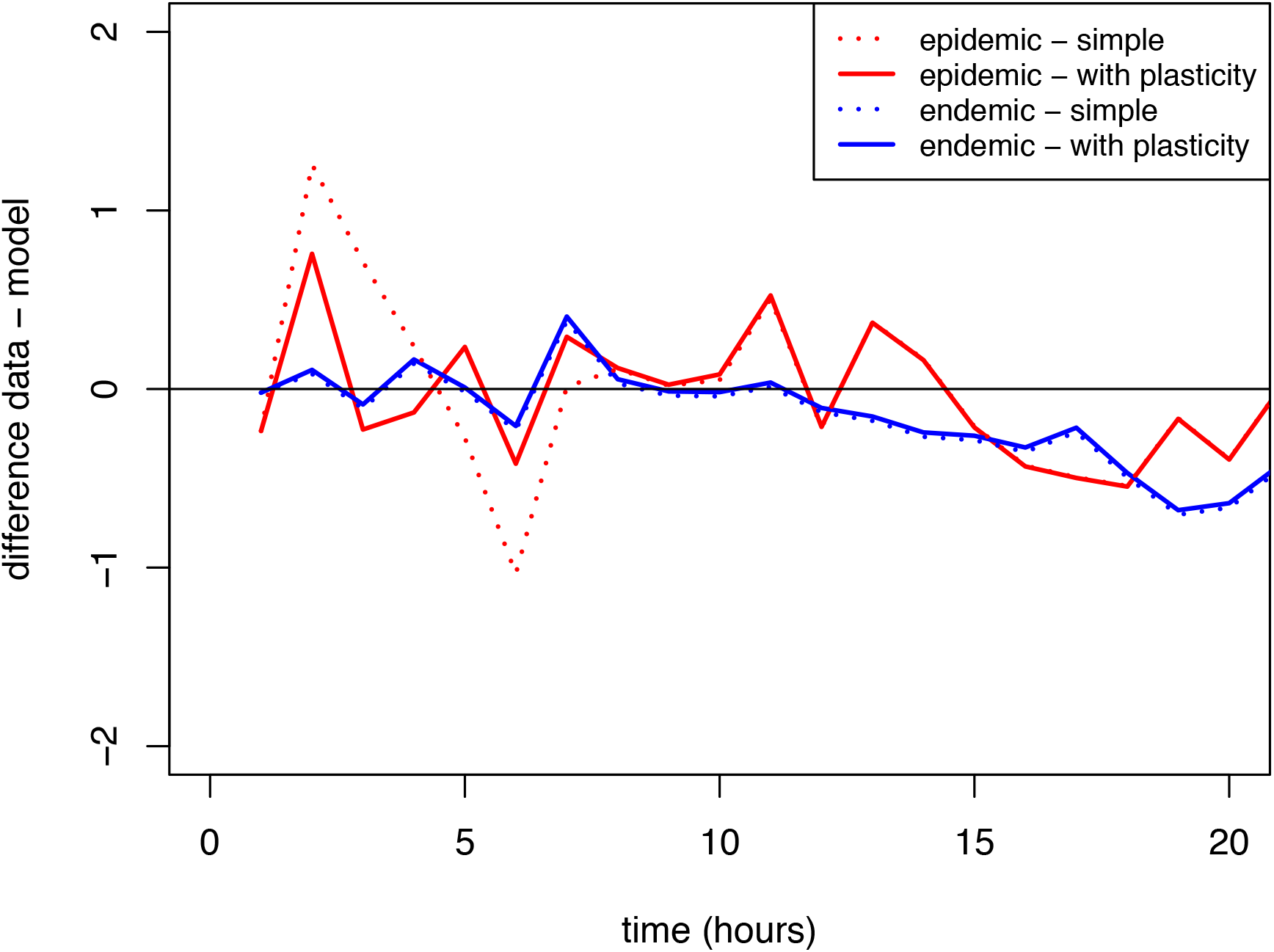
Difference between the model predictions and the data for the simple model without plasticity and the more complex model with plasticity, with a focus on the logit frequency of the mutant free phage, the variable for which most of the difference between the two models arises. The blue curves represent the endemic conditions (high initial prevalence), and the red curve the epidemic conditions (low initial prevalence). The dashed line is the simple model, the plain line is the model with plasticity that explains better the frequency of mutant free phage during the epidemic peak.

